# Working Memory Training Integrates Visual Cortex into Beta-Band Networks in Congenitally Blind Individuals

**DOI:** 10.1101/200121

**Authors:** Johanna M. Rimmele, Helene Gudi-Mindermann, Guido Nolte, Brigitte Röder, Andreas K. Engel

## Abstract

Congenitally blind individuals activate the visual cortex during non-visual tasks. Such crossmodal reorganization is likely associated with changes in large-scale functional connectivity, the spectral characteristics of which can be assessed by analysis of neural oscillations. To test visual cortical integration into working memory networks, we recorded magnetoencephalographic data from congenitally blind and sighted individuals during resting state as well as during a voice-based working memory task prior to and following working memory training with voices, or tactile stimuli or a training-control condition. Auditory training strengthened beta-band (17.5-22.5 Hz) connectivity (imaginary coherency) in the blind and theta-band (2.5-5 Hz) connectivity in the sighted during the task, suggesting different neural coupling mechanisms. In the sighted, theta-band connectivity increased between brain areas involved in auditory working memory (inferior frontal, superior temporal, insular cortex). In the blind, beta-band networks largely emerged during the training, and connectivity increased between brain areas involved in auditory working memory and the visual cortex. The prominent involvement of the right fusiform face area in this beta-band network suggests a task-specific integration of visual cortex. Our findings highlight large-scale interactions as a key mechanism of functional reorganization following congenital blindness, and provide new insights into the spectral characteristics of the mechanism.

## Introduction

The ability to flexibly adapt to a steadily changing environment is a crucial prerequisite for human survival and has been related to the brain’s capacity for structural rewiring and functional reorganization (Buonomano and Merzenich 1998; Pascual-Leone et al. 2005; Röder 2012). In the case of sensory deprivation, such as in congenital blindness, adaptation is required, as individuals have to cope with the lack of information from one of the sensory organs. A large amount of studies suggest that cross-modal plasticity in congenitally blind individuals leads to an increased recruitment of the visual cortex during non-visual tasks (Pascual-Leone et al. 2005; Collignon et al. 2009; Ricciardi and Pietrini 2011; Pavani and Röder 2012), which is often accompanied by higher performance compared to sighted participants (Amedi et al. 2003; Gougoux et al. 2005; Collignon et al. 2009). Importantly, the visual cortex seems not to be merely co-activated but shows task specific involvement (Pietrini et al. 2004; Amedi et al. 2007, 2010; Watkins et al. 2013; Hölig et al. 2014). For example, during tactile or auditory object identity processing (e.g. identifying voices) congenitally blind individuals showed ventral visual stream activation, typically associated with visual feature processing (Pietrini et al. 2004; Amedi et al. 2007, 2010; Gougoux et al. 2009; Hölig et al. 2014). Moreover, a functional relevance of the visual cortex is suggested by transcranial magnetic stimulation studies, as the processing of non-visual tasks was impaired in congenitally blind individuals when activity within the visual cortex was temporarily interfered (Amedi et al. 2003, 2004; Wittenberg et al. 2004; Ptito et al. 2008; Collignon et al. 2009) (for review: Pascual-Leone and Hamilton 2001; Ricciardi and Pietrini 2011).

In a current view, structural and functional brain connectivity strongly determine the functional specification of brain areas (Friederici and Singer 2015; Hannagan et al. 2015). Accordingly, the recruitment of the visual cortex for non-visual tasks in congenitally blind individuals might result in the integration of visual cortex into task-related networks. Studies on changes in structural connectivity following congenital blindness report atrophy of the geniculocortical tracts, while cortico-cortical connections of visual cortex are relatively preserved (Shimony et al. 2006). Although structural connectivity might affect the functional specification of brain areas, direct conclusions about the networks that are involved in a specific task can only be drawn from functional connectivity measures. Some evidence for altered task-related networks in congenitally blind individuals comes from functional magnetic resonance imaging (fMRI) research (Liu et al. 2007; Klinge et al. 2010; Ptito et al. 2012; for review: Lazzouni and Lepore 2014). In line with the assumption that information during non-visual tasks reaches visual cortex through cortico-cortical connections, a recent fMRI study found increased effective connectivity between A1 and V1 in congenitally blind compared to sighted individuals during performance of several auditory tasks (Klinge et al. 2010). These functional connectivity measures, however, might be affected by pre-existing differences between the blind and sighted groups that are not task-related. Furthermore, important information about the spectral characteristics of functional networks cannot be retrieved with the fMRI approach.

Here, we used working memory training to experimentally induce the formation of task-related networks to access the neural mechanisms underlying the integration of visual cortex into working memory networks in congenitally blind individuals. We analyzed changes in networks involved in working memory processing resulting from the training intervention and compared those to changes in matched sighted participants. This approach benefits from the intra-subject paradigm that investigates changes across pre-post sessions, while the within-subject normalization at the same time isolates task-related functional connectivity from pre-existing differences between the groups and provides further insights into the role of visual cortex within working memory related networks. Working memory denotes the ability to temporarily maintain and manipulate information to make it accessible to current cognitive processes (Baddeley and Hitch 1974; Baddeley 2012). The multi-component model (Baddeley and Hitch 1974; Baddeley 2012) suggests that several distinct storage systems are involved in working memory processing (phonological loop, visual sketchpad). Additionally, this model proposes the existence of an executive control and of episodic buffers. Based on findings of largely overlapping activity during memory encoding and maintenance periods, state-based models (Cowan 1995; Oberauer 2002; Esposito and Postle 2015), however, critizised the notion of distinct storage systems. Here, we utilize a central aspect of working memory, that is, its’ capacity limit. Working memory training provides a suitable approach to induce an increase in working memory capacity (for review: Jaeggi et al. 2008; Klingberg 2010; von Bastian and Oberauer 2013). Auditory working memory networks involve ventrolateral prefrontal cortex (Plakke et al. 2015; Cohen et al. 2014; Plakke and Romanski 2014), inferior parietal cortex (IPC) (Jonides et al. 1998; Owen et al. 2005), and the insula (Koelsch et al. 2009; Huang et al. 2013), as well as sensory processing areas (Bancroft et al. 2014; Cohen et al. 2014) (for review: Curtis and D’Esposito 2003; Owen et al. 2005; Plakke and Romanski 2014). Specifically, we used a working memory task where participants had to maintain voice identity information, in order to test whether congenitally blind individuals show a specific integration of brain areas related to visual feature processing, such as regions of the ventral visual stream (Gougoux et al. 2009; Hölig et al. 2014) into the working memory networks.

The dynamics of working memory networks can be investigated by analyzing changes in synchronization of oscillatory brain activity (e.g. using electroencephalography, EEG or magnetoencephalography, MEG), which has been proposed to provide a mechanism for the formation of functional networks (Engel et al. 1992, 2001; Engel and Singer 2001; Jutras and Buffalo 2010; Watrous et al. 2015). Importantly, this approach extends previous fMRI research, as it allows to characterize the spectral characteristics of functional networks, providing additional insights into neural coupling mechanisms involved in crossmodal reorganization (Engel and Fries 2010; Weiss and Mueller 2012; Roux and Uhlhaas 2014; Watrous et al. 2015). In sighted individuals, neural oscillatory activity in the theta- (Roux and Uhlhaas 2014), beta-(Engel and Fries 2010; Kopell et al. 2011; Weiss and Mueller 2012), and gamma-band (Pesaran et al. 2002; Roux et al. 2012) has been associated with working memory maintenance, and theta-and beta-band neural networks have been shown to be modulated by working memory training (Langer et al. 2013; Astle et al. 2015). Previous fMRI studies show activation of fronto-parietal brain areas during working memory processing in congenitally blind individuals, similarly to sighted, plus additional activation of visual cortex (Amedi et al. 2003; Park et al. 2011; Deen et al. 2015). However, very little is known about the neural coupling mechanisms involved in working memory processing in the blind. Coupling mechanisms might be altered in congenitally blind compared to sighted individuals due to changes in the structural and functional architecture of visual areas (Hawellek et al. 2013; Lazzouni and Lepore 2014; Schmiedt et al. 2014). For example, in a recent MEG study, classical resting rhythms in visual cortex have been shown to differ in congenitally blind compared to sighted individuals (Hawellek et al. 2013). Furthermore, research in macaque monkey, using local field potentials (LFPs), suggests that structural changes in visual cortex and thalamo-cortical connections can affect the balance between feedback and feeedforward communication resulting in an altered spectro-temporal profile of neuronal communication (Schmiedt et al. 2014).

A phase-based connectivity measure (imaginary coherency; Nolte et al. 2004) was used to analyze the effects of working memory training with voices on the synchronization of neural activity, recorded with MEG, in the theta-, beta- or gamma-band between brain areas associated with auditory working memory (fronto-parietal, insular and auditory cortex) and visual processing (occipito-temporal cortex) in sighted and congenitally blind individuals. An advantage of using imaginary coherency as connectivity measure is that it is insensitive to spurious brain connectivity that can result from volume conduction. We provide electrophysiological evidence that in sighted participants working memory training results in a strengthening of theta-band connectivity across brain areas associated with auditory working memory, while in the congenitally blind beta-band connectivity increased between brain areas associated with auditory working memory and visual cortex. Our results indicate a training-induced integration of visual cortex into working memory networks in the blind and suggest that the spectral characteristics of networks associated with working memory training differ in blind and sighted individuals.

## Materials and Methods

### Experimental Paradigm

The experiment was part of a larger working memory training study (Table S1) (the remaining data will be reported elsewhere).

#### Pre-and post-training sessions: 2-back task

The MEG data were recorded in two sessions: a pre-training session preceding and a post-training session following extensive working memory training. In an auditory 2-back task participants were instructed to continuously indicate whether the current speaker (10 different speakers) of a pseudo-word matched the speaker of the next-to-last stimulus (Fig. 1 A). Congenitally blind and sighted participants were assigned to three training conditions: (1) working memory training with voices; (2) working memory training with tactile motion stimuli;(3) an active training-control task. Additionally, we here report data that were recorded during resting state (∼ 3.5 min) prior to the 2-back task in the pre-training session.

**Figure 1:**
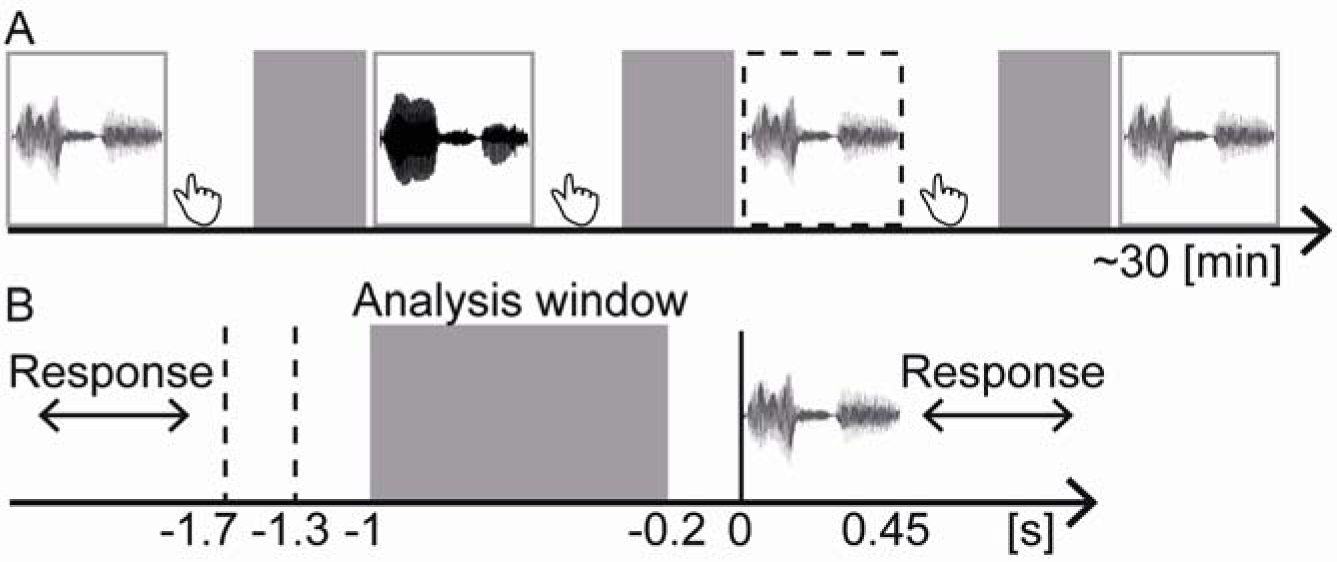
Schematic of the Voice Recognition Task. (A) In an auditory 2-back task the pseudo-word “befa” (waveform) pronounced by 10 different speakers (speaker identity is color coded) was presented. Participants had to indicate by button press whether the current speaker matches the speaker presented two stimuli ago. A target (red frame) required a “yes” response, while a non-target (grey frame) required a “no” response. The duration of the experiment depended on the response times (∼ 30 min). (B) MEG activity was analyzed in the analysis window (−1 to −0.2 pre-stimulus) during the delay period. During the delay period (A, B: grey boxes) participants were maintaining items in memory. Each response was followed by an inter-trial interval (ITI) that was randomly jittered between 1300-1700 ms. The minimum ITI lasted from −1300 ms to stimulus onset (0 ms) in all participants. To avoid overlap with response- and stimulus-related processing the first 300-700 ms and last 200 ms of this ITI were discarded for the analyses.

Throughout the MEG recording, participants were seated in a comfortable chair. All participants were blindfolded. Instructions were provided orally to each participant in the beginning of the experiment. Throughout the experiment instructions and auditory stimuli were presented through insert ear-plugs (E-A-RTONE Gold 3A Insert Earphones, Ulrich Keller Medizin-Technik, Weinheim, Germany) at normal conversational level (∼75 dB SPL). Prior to the experiment all participants performed several blocks of the working memory task to ensure task comprehension (an error sound provided direct feedback). An extended version of the “Verbaler Lern-und Merkfähigkeitstest” (VLMT; German version of the Auditory Verbal Learning Test) was conducted after the pre- and post-training recording session. The VLMT was extended by instructing participants to remember the items in the correct order if possible, and additionally analyzing scores for absolute (counting correct responses up to the first error) and relative (counting all responses where a correct sequence of minimum two items was reported) temporal order memory. Following Jaeggi et al. (2007), we accessed participants working memory strategies using an inventory that was conducted after the last experimental session. Participants indicated whether they used any of the following strategies: verbal memorization, internal rehearsal, internal sequencing, story telling, visual imaging, spatial imaging, episodic memorizing, whether they performed the task intuitively, or whether they had no strategy (cf. supporting information).

Auditory stimuli consisted of a pseudo-word (“befa”), with a stimulus length of 450ms, spoken by 10 different speakers (5 female). All stimuli were peak normalized. We used pseudo-words to avoid effects related to speech semantics, such as semantic associations, or word familiarity, which might affect the processing of voice identity differently in the blind and sighted participants. Participants were instructed to prioritize response accuracy over speed. Each response was followed by an inter-trial interval that was randomly jittered between 1300-1700 ms. The 2-back task comprised 15 blocks in the pre-training session, and 12 blocks in the post-training session with 32 (30+n) stimuli per block. For each block, stimulus sequences were quasi-randomly designed using Matlab: (1) positions of targets (30%) and non-targets were randomized; (2) stimuli were quasi-randomly assigned as targets or non-targets; (3) 19% catch trials (i.e., 1-back and 3-back stimulus pairs) were randomly inserted.

#### Training and training-control sessions: n-back task

Congenitally blind and sighted participants of the different training conditions were matched with respect to age, gender, and handedness (Table 1). The training comprised four sessions (∼ 2-3 hours duration each). During the working memory training with voices participants performed an adaptive auditory n-back task and during the working memory training with tactile motion stimuli an adaptive tactile n-back task (30 blocks of 30+n stimuli). In the auditory n-back task, participants had to indicate whether the current speaker matches the n-to-last speaker. In the tactile n-back task, participants had to indicate whether the current motion direction and stimulated finger match those of the n-to-last stimulus. A programmable mechanical Braille stimulator (QuaeroSys Medical Devices, Schotten, Germany) with 5 stimulator units (2x4 pin matrix) was used to generate an up- or a down movement at each of the five fingers of one hand (the stimulated hand was matched between groups and conditions). This resulted in 10 tactile apparent motion stimuli. The first training session started with a 2-back block. The HR and correct rejection rate (CRR) were used to adapt the task demands to the individual performance:

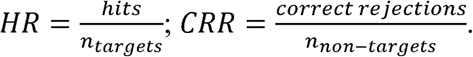

**Table 1:**
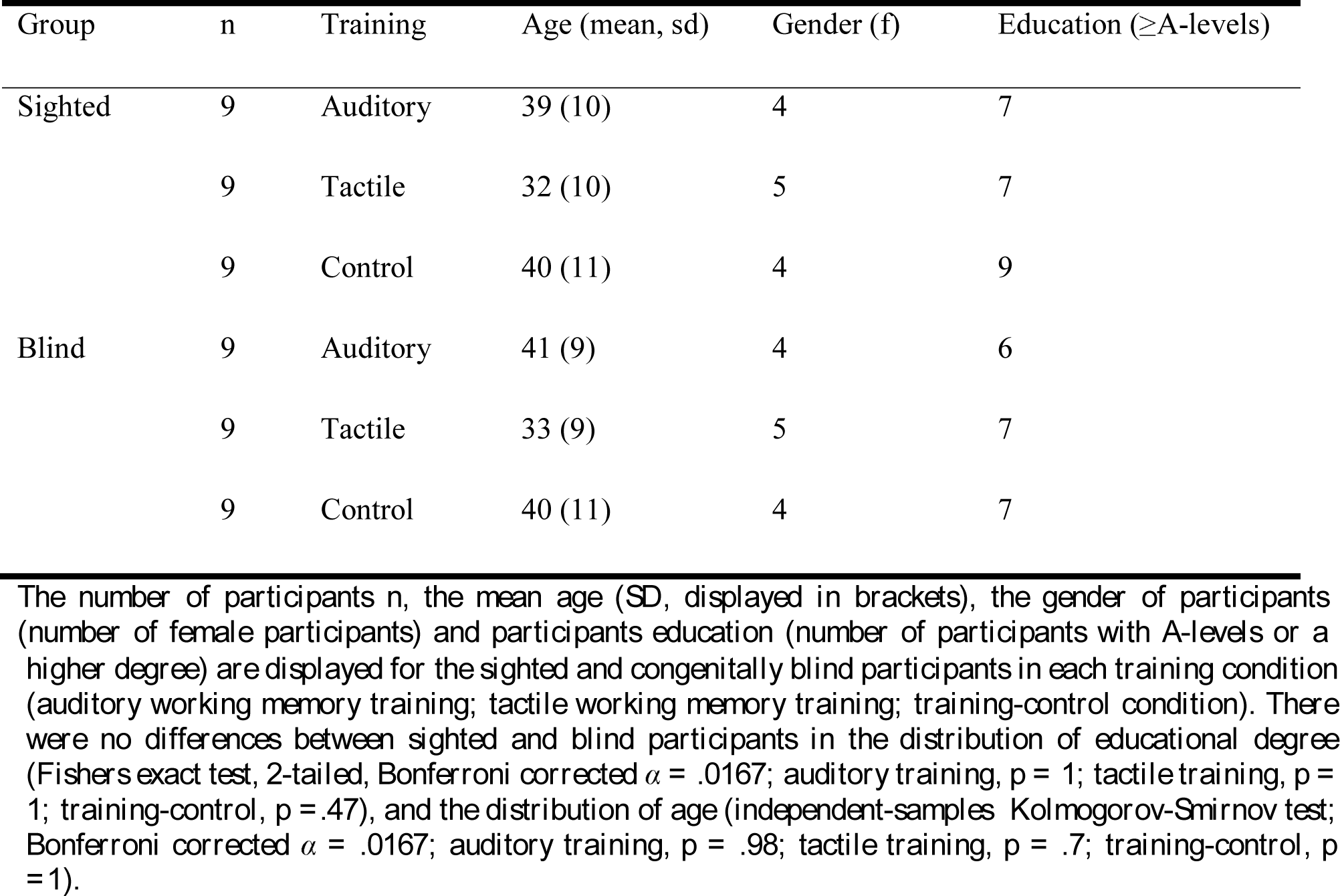
Sample Demographics

| Group | n | Training | Age (mean, sd) | Gender (f) | Education (≥A-levels) |
| --- | --- | --- | --- | --- | --- |
| Sighted | 9 | Auditory | 39 (10) | 4 | 7 |
|  | 9 | Tactile | 32 (10) | 5 | 7 |
|  | 9 | Control | 40 (11) | 4 | 9 |
| Blind | 9 | Auditory | 41 (9) | 4 | 6 |
|  | 9 | Tactile | 33 (9) | 5 | 7 |
|  | 9 | Control | 40 (11) | 4 | 7 |

*The number of participants n, the mean age (SD, displayed in brackets), the gender of participants (number of female participants) and participants education (number of participants with A-levels or a*
*higher degree) are displayed for the sighted and congenitally blind participants in each training condition (auditory working memory training; tactile working memory training; training-control condition). There were no differences between sighted and blind participants in the distribution of educational degree (Fishers exact test, 2-tailed, Bonferroni corrected $\alpha = .0167$ ; auditory training, $p = 1$ ; tactile training, $p = 1$ ; training-control, $p = .47$ ), and the distribution of age (independent-samples Kolmogorov-Smirnov test; Bonferroni corrected $\alpha = .0167$ ; auditory training, $p = .98$ ; tactile training, $p = .7$ ; training-control, $p = 1$ ).*

When participants showed high performance in a block, the n was increased (HR ≥ 70 % and CRR ≥ 75%). The n was decreased after low performance (HR < 60 % or CRR < 60 %). Otherwise the n did not change. Participants in the training-control condition conducted a 1-back task with low cognitive demand in both modalities. The stimuli for the 1-back task were the same as used in the auditory and tactile n-back task, respectively. Participants indicated whether the current stimulus matched the last stimulus. The auditory and tactile 1-back task were performed on alternating training days, while we balanced which modality was tested first (overall 40 blocks á 30+n stimuli; note that more blocks were required for the easy 1-back compared to the n-back task to achieve the same total training duration of approximately 8 hours per participant. Note that the training duration in one single modality was reduced in the training-control compared to the training conditions.). The active control condition was chosen to differentiate working memory training effects from more general effects, such as repetition or intervention effects. Note that we investigated the elaboration of brain networks as a consequence of training. Auditory and tactile working memory tasks were selected that allowed for improvements in all individuals. By analyzing the neural correlates of individual training effects, we controlled for possible differences in pre-training skills in auditory and tactile processing.

### Subjects

Congenitally blind (n = 27) and sighted (n = 27) matched control participants took part in the study. All participants were healthy with normal hearing (self-report) and no history of psychiatric or neurological disorders. One blind participant reported a history of depressive mood disorder, however, with no current symptoms or need for treatment. Sighted participants had normal or corrected to normal vision (self-report). Due to loss of vision, following (pre)natal anomalies in the peripheral visual system (retinopathy of prematurity: n = 9; genetic defect, n = 5; congenital optic atrophy: n = 3; Leber′s congenital amaurosis: n = 2; congenital cataracts, glaucoma: n = 2; congenital retinitis: n = 2; binocular anophthalmia: n = 2; retinitis pigmentosa: n= 1; congenital degeneration of the retina, n = 1), congenitally blind participants were totally blind since birth with minimal residual light perception in 18 participants. During the training sessions all participants underwent additional psychological tests, including the German version of the PSQ-20 (Perceived Stress Questionnaire, Levenstein et al. 1993; German modified version, Fliege et al. 2005); 2); the HSWBS (German Habituelle Subjektive Wohlbefindens Skala; Dalbert 1992) to measure wellbeing; and the MWT-B (German Mehrfachwahl-Wortschatz-Test, Lehrl 2005) to assess verbal intelligence. The blind and sighted participants did not differ in any of the assessed psychological variables (PSQ-R20: t(45.06) = .38, *p* = .704; HSWBS: t(46.55) = .46, *p* = .647; MWT-B: t(48.14) = .67, *p* = .507).

The study was approved by the German Psychological Association (DGPs). All participants gave written informed consent prior to the experiments and received monetary compensation for participation.

### Behavioral Data Analysis

For the behavioral data analysis all responses within a response window (2-6 sec) were analyzed. The first two trials of each block were removed (participants were informed that a “non-target response” was correct for these initial trails). The hit rate (HR) was calculated separately for each participant and session (pre, post). Analyses of Variance (ANOVAs) were used to test training-related differences in working memory performance increases across sessions (post minus pre HR) with the between-subject factors training condition (auditory, tactile, control) and group (blind, sighted).

### MRI and MEG Data Acquisition

T1-weighted structural MRI scans were obtained for each participant except for those who did not meet the MRI scan criteria (sighted: n = 3). The MRI recording was performed on a 3 Tesla scanner (Siemens Magnetom Trio, Siemens, Erlangen, Germany). The MEG data were recorded in a magnetically shielded room using a 275-channel whole-head system (Omega 2000, CTF Systems Inc.). The electrooculogram and electrocardiogram were derived for offline artifact rejection. Prior to and after each experiment the head position was measured relative to the MEG sensors. The head position was tracked during the recording and head displacement was corrected in the breaks using the fieldtrip toolbox (http://fieldtrip.fcdonders.nl; Stolk et al. 2013). The data were recorded with a sampling rate of 1200 Hz and low pass filtered online (cut-off: 300 Hz).

### MRI Data Analysis

The FieldTrip toolbox (http://fieldtrip.fcdonders.nl; (Oostenveld et al. 2011) and other customized Matlab toolboxes were used for data analyses. Anatomical landmarks (nasion, left and right pre-auricular points) were manually identified in the individual MRIs. Individual MRIs of all participants were then analyzed by, first, obtaining probabilistic tissue maps (including cerebrospinal fluid white and gray matter) from the anatomical MRI. The standard Montreal Neurological Institute (MNI) brain was used for cases without an individual MRI. Second, a single shell volume conduction model (Nolte 2003) was used to estimate the physical relation between sensors and sources. A source model was generated using a regular 3-D grid (0.8 cm spacing). The grid was used to generate a warped MNI grid to project individual MRIs on the standard MNI brain. Finally, based on the warped MNI grid and the probabilistic tissue map a leadfield (forward model) was calculated that was used for source reconstruction.

### MEG Data Analysis

For preprocessing, the data were band-pass filtered off-line (1–160 Hz, Butterworth filter; filter order 4) and line-noise was removed using bandstop filters (49.5-50.5, 99.5-100.5, 149.5-150.5 Hz, two-pass; filter order 4). Only correct responses and, matched with the behavioral data analyses, only trials where participants responded within a 2-6 sec window were analyzed, while the first two trials of each block were removed. In a semi-automatic artifact detection procedure, trials containing muscle artifacts, signal jumps, and slow artifacts were rejected. The continuous data were epoched with respect to stimulus onset (-2 to 2 sec) and down sampled to 500 Hz. Independent component analysis (infomax algorithm; Makeig et al. 1996) was used to remove eye-blink, eye-movement and heartbeat-related artifacts based on the component topography, time course and variance across trials (components were first reduced to 64 components using principal component analysis). The number of trials was matched across pre- and post-training sessions for each participant by randomly removing trials from the larger set. This resulted in the following amount of trials in each session (pre, post) for the sighted participants: Auditory training, mean = 319 (sd = 15), tactile training, mean = 307 (sd = 26), training-control, mean = 292 (sd = 61); and for the blind participants: auditory training, mean = 286 (sd = 33), tactile training, mean = 297 (sd = 26), training-control, mean = 293 (sd = 34); overall sighted participants, mean = 306 (sd =40); overall blind participants, mean = 291.8 (sd = 30). Data in the delay window (-1 to – 0.2 s pre-stimulus; 0.8 s segments) were further analyzed. The data recorded during resting state were processed similarly. The continuous data were epoched into 0.8 s segments to match the auditory 2-back data. The number of trials was matched between the pre-training auditory 2-back data and the resting state data separately for each participant: Sighted participants, mean = 298 (sd = 41); congenitally blind participants, mean = 293 (sd = 37).

### Source Power and Connectivity Analysis

Discrete Fourier transform was performed at 2.5-100 Hz, separately for each participant and session (pre, post, resting state) (segment length: 0.4 s; segment shift: 0.05 s; frequency resolution: 2.5 Hz). The cross-spectrum was retrieved by calculating the cross-covariance function of the Fourier transformed data (Nolte et al. 2004). Given that x_i_(f) and x_j_(f) are the Fourier transforms of the time series x_i_(t) and x_j_(t) of sensor i and j the cross-spectrum (cs) was calculated as:

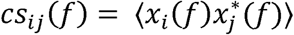

Exact Low-Resolution Brain Electromagnetic Tomography (eLoreta) was used to calculate a spatial filter based on the individual leadfield (2982 inside brain voxel, 3 dipole directions) (Pascual-Marqui 2007).

#### Source Connectivity Analysis

Based on the automated anatomical labeling (AAL) procedure (Tzourio-Mazoyer et al. 2002) five regions of interest (ROIs) were selected: frontal, parietal, insula, temporal and occipito-temporal; Additionally, ROIs in V1 and V2 were selected, and the Brede database was used to select coordinates of the left and right fusiform face area (FFA) and medial temporal areas (hMT+/V5) (Table S2).

The cross-spectrum was averaged across frequencies for three frequency bands: theta (2.5-5 Hz), beta (17.5-22.5 Hz) and gamma (40-60 Hz). For each frequency and each pair of signals imaginary coherency was calculated as the complex cross-spectrum normalized by the square root of the product of powers. For each pair of voxels dipole directions were estimated by maximizing imaginary coherency values (Ewald et al. 2012). Fischer’s z-transformation was used for variance stabilization (inverse hyperbolic tangent). In order to estimate imaginary coherency between each voxel and a ROI, we averaged the connectivity between that voxel and all voxels of an AAL brain region (Table S2), first, per hemisphere and, second, across hemispheres and across all AAL regions of a ROI. The imaginary coherency during resting state, and the differences between sessions (auditory 2-back post-minus-pre; auditory 2-back pre-minus-resting state) were further analyzed. The (maximized) imaginary coherency values range between 0 and 1, with 1 indicating maximally synchronous time series.

#### Source Power Analysis

For each voxel, source power (pow) was calculated based on the individual Nx3 spatial filter (*A*), for N channels and 3 dipole directions, and cross-spectra (cs) at each frequency (3-100 Hz) separately (Pascual-Marqui 2007).

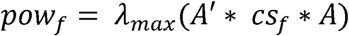

where **λ**_max_ denotes the maximum eigenvalue. Source power was averaged for each participant and each session and averaged across frequencies of each frequency band (theta: 2.5-5 Hz, beta: 17.5-22.5 Hz, gamma: 40-60 Hz). Source power (resting state) and source power contrasts between sessions were further analyzed: 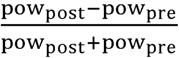 and 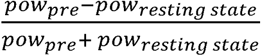

### Permutation Statistics

Independent-sample random permutation statistics were performed to test connectivity and power changes in the auditory 2-back task separately for the three frequency bands (theta, beta, gamma). Only effects with a minimum of 100 voxels with significant differences are reported. In all analyses, false discovery rate (FDR) (Genovese et al. 2002) with Q values equal 0.2 was used to control for multiple comparisons of voxels, ROIs and frequency bands, if not stated otherwise. In case no solution was found with the FDR correction, the Bonferroni correction was automatically applied (Q/n).

First, we tested differences in training-related connectivity and power changes between the blind and sighted participants. Training-related changes (i.e., differences in connectivity or power changes across sessions, using the individual post-minus-pre difference for connectivity and the contrast for power, between the working memory training with voices and the training-control condition) in the blind were tested against training-related changes in the sighted participants. Differences were tested by randomly permuting participants’ group affiliation (blind, sighted) (10,000 permutations) separately for the auditory training and training-control condition. Training-related changes were then calculated, (1), separately for participants of each group and, (2) for the randomly permuted groups. Finally, we tested the observed group difference (blind minus sighted) against the differences of the randomly permuted groups.

Second, effects of working memory training on connectivity and power changes across sessions (using the individual post-minus-pre difference for connectivity and the contrast for power) during the auditory 2-back task were tested separately for the blind and sighted participants. We tested differences in changes across sessions between the training conditions by randomly permuting participants’ training condition affiliation (10,000 permutations) and comparing the observed condition difference against the difference of the randomly permuted conditions.

The resting state data of one participant had to be discarded due to file distortion resulting in the analysis of data from 26 congenitally blind and 27 sighted participants for the analysis of the pre-training data. In an exploratory analysis, where we only controlled for multiple comparisons of voxels, we tested whether differences in the working memory networks of blind and sighted participants already existed prior to the training. Independent-sample permutation statistics were performed to test differences between blind and sighted participants during the auditory 2-back task and during resting state prior to the training. The analyses were performed on the difference of the connectivity data recorded during the pre-training auditory 2-back task and resting state, as well as on the contrast of the power data recorded during the pre-training auditory 2-back task and resting state. Participants’ group affiliations (blind, sighted) were randomly permuted (10,000 permutations). The observed group differences were tested against the differences of the randomly permuted groups. Furthermore, we similarly analyzed the pre-training connectivity and power data recorded during the resting state to access baseline differences in power and connectivity between blind and sighted participants.

## Results

### Working Memory Training with Voices Increases Performance

Participants’ high pre-training performance (HR sighted: auditory training, mean = .87, sd = .09; tactile training, mean = .91, sd = .06; training-control, mean = .87; sd = .08; HR blind: auditory training, mean = .77, sd = .13; tactile training, mean = .89, sd = .09; training-control, mean = .91; sd = .05) shows that participants were able to perform the task. As expected, working memory training with voices resulted in a HR increase in the 2-back task with voices (Fig. S1; main effectη pof condition, F(2,48) = 4.96, *p* < .0500, 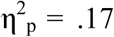). Participants who performed the working memory training with voices showed a higher HR increase across sessions compared to those in the training-control condition (post-hoc independent-sample Student’s t-tests; p = .0123; Bonferroni corrected α:. 0167) and to those who underwent tactile working memory training (p =.0161). In contrast, working memory training with tactile stimuli did not improve performance compared to the training-control condition (p = .5939) indicating a modality-specific training effect. No differences in working memory training effects were observed between blind and sighted participants (no main effect of group; p = .5453; and no interaction of group and condition; p = .7174). As an adaptive working memory training with voices, but not with tactile motion stimuli, compared to a training-control condition resulted in a performance increase in the auditory 2-back task, for the analyses of the neurophysiological data the effects of the adaptive working memory training with voices on working memory networks were compared to those of the training-control condition (for data on the tactile training condition: Fig. S5 B).

### Training Integrates Visual Cortex into a Beta-Band Network in the Congenitally Blind

To test effects of working memory training on visual cortex integration into working memory networks, we analyzed connectivity and power during the delay period where participants were maintaining items in memory during the 2-back task with voices (Fig. 1B). Differences in training-related connectivity and power changes between participants with working memory training with voices and those in the training-control condition were analyzed in the theta-, beta- and gamma-bands and compared between blind and sighted participants. Connectivity was analyzed for ROIs related to auditory working memory (frontal, parietal, insula, temporal) and visual processing (occipito-temporal; Table S2).

The blind participants showed a stronger training-related increase in beta-band connectivity compared to the sighted between visual cortex and brain areas associated with auditory working memory (Fig. 2A and Fig. S5 A; Q = 0.2; FDR corrected α = .0079; *p*-values < .0079). Connectivity particularly increased between the frontal, insula and temporal ROIs and occipito-temporal brain areas (including V2, the right fusiform gyrus, and the right inferior temporal lobe, ITL); between the frontal ROI and right inferior parietal gyrus (IPG); and between the occipito-temporal ROI and the inferior frontal gyrus (IFG), the middle frontal gyrus (MFG), the right superior temporal gyrus (STG), the anterior superior temporal sulcus (STS), and the right insula.

**Figure 2:**
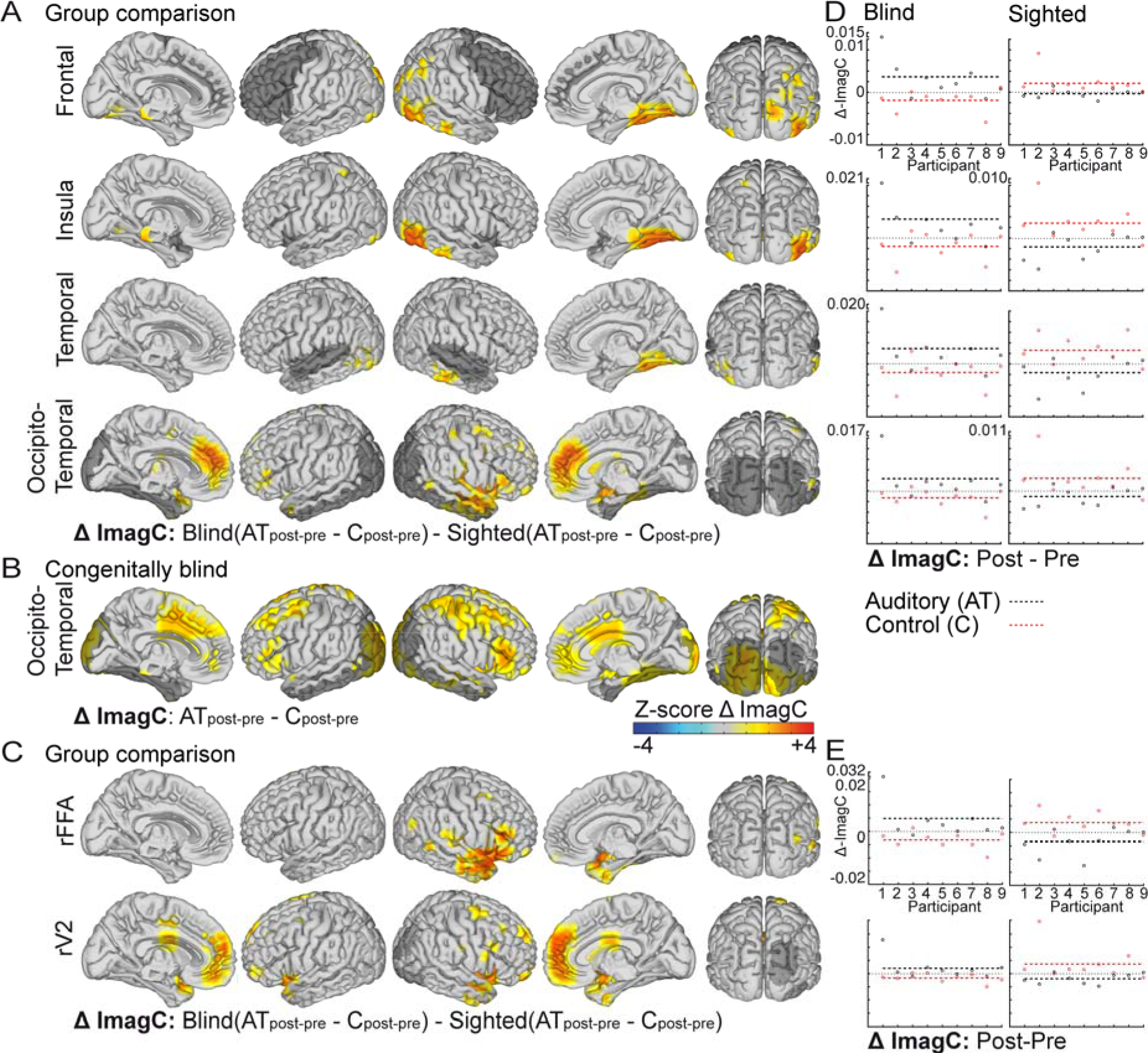
Increased Beta-Band Connectivity in the Congenitally Blind. *(A) Congenitally blind participants showed training-related (AT, auditory training; C, control condition) increases in connectivity in the beta-band compared to the sighted between the frontal, insula, temporal and occipito-temporal regions of interest (ROIs; displayed in dark transparent gray) and the voxels displayed in color. Each row shows effects for the ROI labeled on the left. Higher connectivity values in the blind compared to the sighted participants are displayed in warm colors. (B) In blind participants working memory training with voices increased connectivity across sessions in the beta-band between the occipito-temporal ROI (displayed in dark transparent gray) and the voxels displayed in color compared to the training-control condition. Higher connectivity increases in the working memory training with voices compared to the training-control condition are displayed in warm colors. (C) Congenitally blind participants showed increased training-related connectivity in the beta-band compared to the sighted between the right fusiform face area (FFA) ROI (displayed in dark transparent gray) and the voxels displayed in color and between the right V2 ROI (displayed in dark transparent gray) and the voxels displayed in color. Higher connectivity values in the blind compared to the sighted participants are displayed in warm colors. In (A-C) connectivity differences are displayed as z-scores (connectivity differences divided by the SD of the permutations). The distribution of participants′ connectivity values is plotted on the right for the main comparison (D) and the comparison with specific ROIs in the ventral and dorsal visual stream (E). The post-minus-pre difference in connectivity (imagC) at each ROI, averaged across voxels with significant effects, is plotted separately for the auditory training condition (black circles) and the control condition (red circles), for the sighted and blind participants respectively. The mean connectivity difference of each training condition is displayed as dashed line. Note that several data points are plottet offset with the value displayed at the y-axis.*

An analysis of theta-, beta-and gamma-band connectivity, where we controlled for multiple comparisons of voxels, ROIs and frequency-bands, in the congenitally blind revealed no training-related connectivity changes across sessions. However, in order to further explore the training-related connectivity differences between the sighted and the blind, we performed an exploratory analysis of the theta-and beta-band with a more liberal control for multiple-comparison (multiple comparisons of voxels at each ROI while not for the amount of ROIs and frequency bands).

We confirmed that the observed differences between sighted and blind participants in the beta-band originated from training-related connectivity increases in the blind (Fig. 2B; Fig. 3). Blind participants with working memory training showed stronger beta-band connectivity increases across sessions compared to those in the control-condition at the occipito-temporal ROI (Q = 0.2; FDR corrected α = .0531; p-values < .0531; Fig. 2B). Connectivity increased between the occipito-temporal ROI and the IFG, the MFG, the pre-and post-central gyrus, and parts of occipital cortex. Working memory training did not affect connectivity in blind participants at the beta-band at any other ROI (Q = 0.2; at all other ROIs FDR corrected α = .0001; auditory: p-values ≥ .0011; frontal, insula, parietal: p-values ≥ .0004), or the theta-band (at all ROIs FDR corrected α = .0001; auditory: p-values ≥ .0012; occipito-temporal: p-values ≥ .0106; frontal: p-values ≥ .0110; insula, parietal: p-values ≥ .0051).

**Figure 3:**
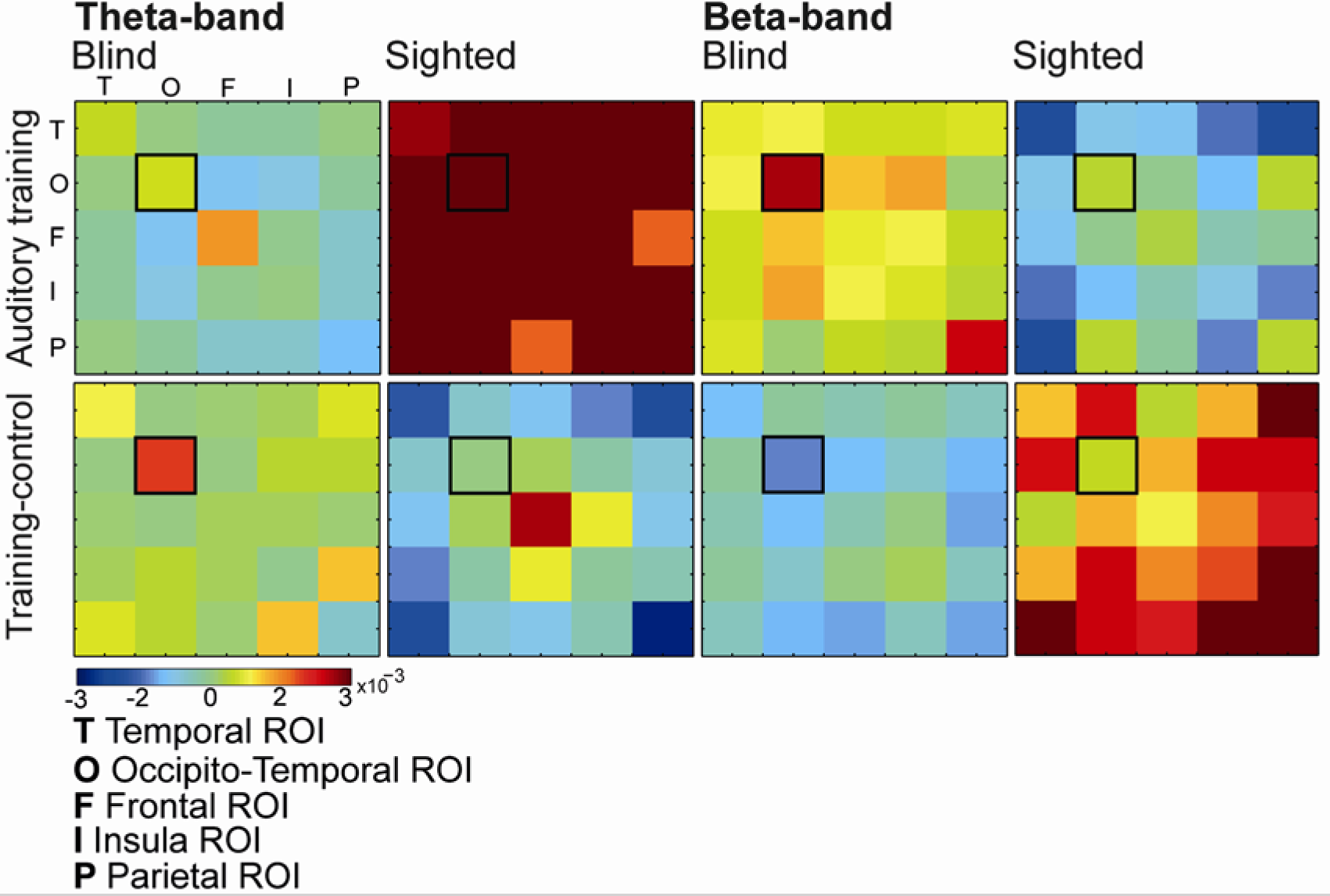
Connectivity Matrix. *The difference in connectivity (imaginary coherency) across sessions (post-minus-pre) between all ROIs (temporal, occipito-temporal, frontal, insula, parietal) is displayed separately for the sighted and congenitally blind participants, the auditory working memory and training-control condition and the theta and beta frequency band. Connectivity was averaged separately for each participant, for each voxel between that voxel and all voxels of each ROI, as described in the Materials and Methods section. This resulted in connectivity measures between each ROI and all voxels. For the purpose of illustration ROI-ROI connectivity pairs were computed, by additionally averaging connectivity for each ROI between that ROI and all voxels of each of the five ROIs. The figure illustrates the connectivity patterns in the different groups and training conditions, particularly, the increase in theta-band connectivity in the sighted participants with auditory training, and in beta-band connectivity in the blind participants with auditory training. Connectivity within occipito-temporal ROIs is highlighted (black box). Interestingly an antagonism between theta-and beta-band networks can be observed, such that when working memory training resulted in an increased connectivity in one frequency band, it rather decreased in the control group while connectivity in the other frequency band showed the opposite pattern across training groups (cf. discussion section). Note the figure does not refer directly to the statistical analyses.*

In sum, working memory training with voices resulted in differences between congenitally blind and sighted participants in the beta-band. Further analyses showed working memory training increased beta-band connectivity between visual areas and an auditory working memory network in the blind.

### Beta-Band Connectivity of the Fusiform Face Area Increases in the Blind

In previous brain imaging studies, congenitally blind individuals showed task-specific activation of visual cortex (Hölig et al. 2014), whereby the right fusiform face area (FFA) was activated during voice identity processing. In the present study, blind participants showed a stronger training-related increase in beta-band connectivity across sessions compared to the sighted between the right FFA ROI and the right IFG, STG and insula (Fig. 2C; Q = 0.2; FDR corrected α = .0031; all p-values < .0031), and between the right V2 ROI and the insula, the right STS/STG and frontal brain areas. There were no effects at the left FFA ROI, the dorsal visual stream ROIs (left and right hMT+/V5), and the left and right V1 ROIs (p-values > .0031). There were no effects in the theta-band (p-values > .0031). The findings suggest that training in a n-back task with voices results in the extension of auditory working memory networks in the congenitally blind compared to the sighted, particularly, by integrating visual brain regions associated with face processing in sighted individuals.

### Training Strengthens a Theta-Band Network in the Sighted

The sighted participants showed a stronger training-related increase in theta-band connectivity compared to the blind between brain areas associated with auditory working memory. The training-related increase in connectivity in the sighted compared to the blind was observed at the frontal, parietal, insula and temporal ROIs (Fig. 4A and Fig. S5 A; Q = 0.2; FDR corrected α =.0079; p-values < .0079). Connectivity differences were most pronounced between the frontal ROI and the right insula, the left STG, the left inferior parietal lobe (IPL), the anterior middle temporal gyrus (MTG), and parts of left V1; between the parietal ROI and the left IFG and MFG, and the anterior MTG; between the insula ROI and right MTL, the left IPG the left posterior STG and STS, and left V1; and between the temporal ROI and IFG, and anterior MTG. There were no effects at the occipito-temporal ROI (p-values ≥ .0081).

**Figure 4:**
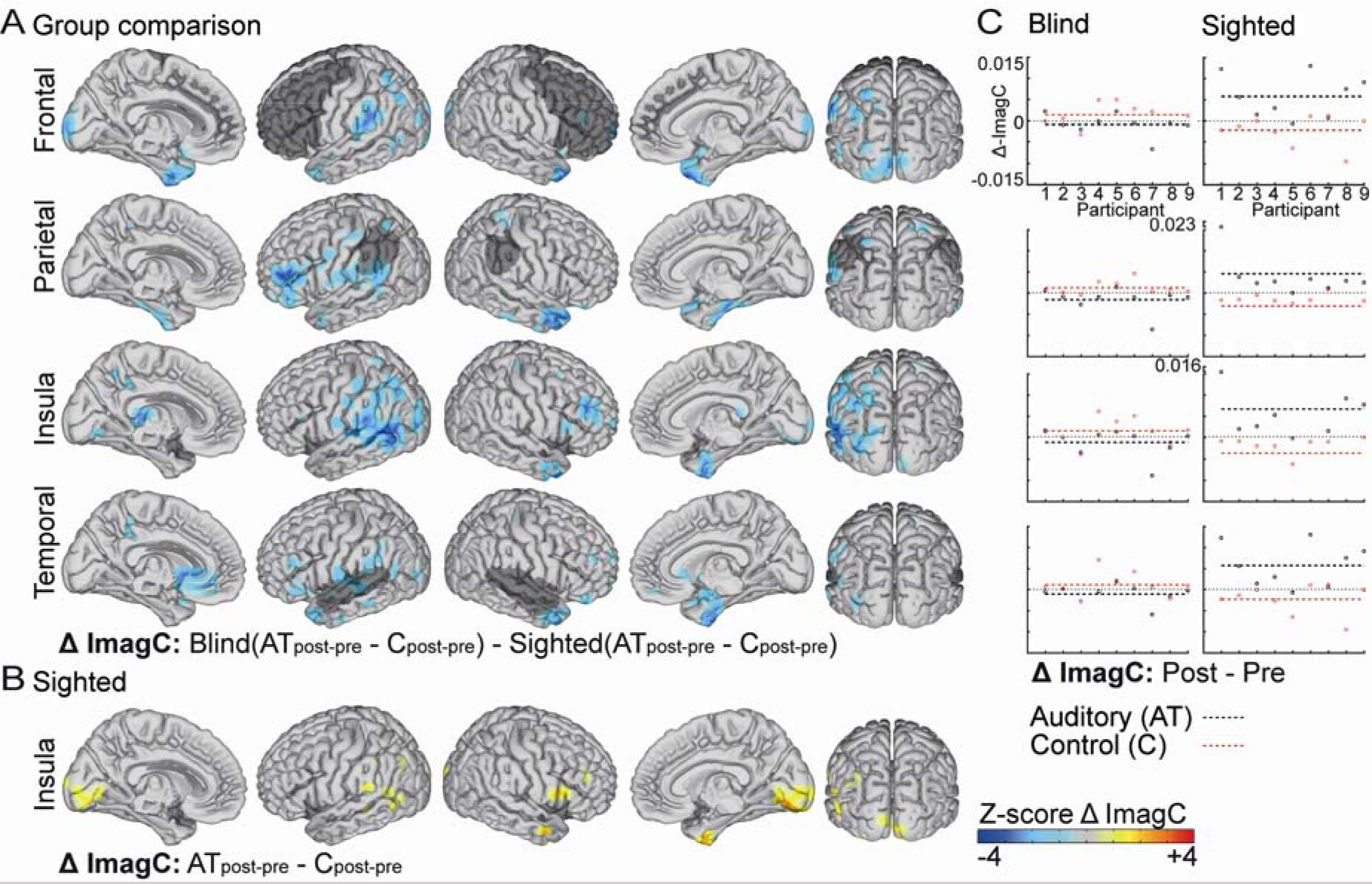
Increased Theta-Band Connectivity in the Sighted. *(A) Sighted participants showed increased training-related connectivity (AT, auditory training; C, control condition) in the theta-band compared to the congenitally blind between the frontal, parietal, insula, and temporal ROIs (displayed in dark transparent gray) and the voxels displayed in color. Each row shows effects for the ROI labeled on the left. Note that cold colors highlight connectivity increases in the sighted compared to the congenitally blind. (B) In sighted participants working memory training with voices increased connectivity across sessions in the theta-band between the insula ROI (displayed in dark transparent gray; labeled on the left) and the voxels displayed in color compared to the training-control condition. Higher connectivity increases in the working memory training with voices compared to the training-control condition are displayed in red. In (A-B) connectivity differences are displayed as z-scores. (C) The distribution of participants connectivity values is plotted on the right. The post-minus-pre difference in connectivity (imagC) at each ROI, averaged across voxels with significant effect, is plotted separately for the auditory training condition (black circles) and the control condition (red circles), for the sighted and blind participants respectively. The mean connectivity difference of each training condition is displayed as dashed line. Note that several data points are plotted offset with the value displayed at the y-axis.*

In an analysis of theta-, beta- and gamma-band connectivity in the sighted we confirmed that the observed changes were due to training-related connectivity increases across sessions in the sighted participants. The sighted participants with working memory training with voices showed a stronger increase in theta-band connectivity across sessions compared to participants in the training-control condition (Fig. 4B; Q = 0.2; FDR corrected α = .0018; all *p*-values < .0018; Fig. 3) between the insula ROI and the right anterior MTG, the right IFG, the left STG, and parts of visual cortex. In the sighted participants, there were no significant differences at any other ROI (frontal: p-values ≥ .0019; parietal: p-values ≥ .0019; temporal: p-values ≥ .0019; occipito-temporal: p-values ≥ .0019), or in beta- and gamma-band (beta-band: frontal: p-values ≥ .0020; insula: p-values ≥ .0020; parietal: p-values ≥ .0022; temporal: p-values ≥ .0025; occipito-temporal: p-values ≥ .0020; gamma-band: frontal: p-values ≥ .0787; insula: p-values ≥ .0403; parietal: p-values ≥ .0343; temporal: p-values ≥ .0532; occipito-temporal: p-values ≥ .1669) connectivity across sessions between participants of the auditory working memory training vs. control condition.

In sum, working memory training with voices strengthened a theta-band auditory working memory network in the sighted.

### Local Activity was not Affected by Working Memory Training

Besides large-scale connectivity, working memory training with voices might impact local activity within areas relevant for working memory processing differently in blind and sighted participants. However, we did not find working memory training-related power differences between the congenitally blind and sighted at any frequency band (Q = 0.2; all frequency bands FDR corrected α = .0001; theta-band: all p-values ≥ .0012; beta-band: all p-values ≥ .0061; gamma-band: all p-values ≥ .0002), and no working memory training effects on power in any group (Q = 0.2) (for a ROI specific analysis: Fig. S4 A, Fig. S4 B). An inspection of the overall power-spectrum (averaged across voxels, pre and post training sessions and training conditions) shows reduced alpha-band power in the congenitally blind compared to the sighted participants (Fig. S2 A). Reduced alpha-band power in the blind across visual cortex is a well-known finding (Noebels et al. 1978; Kriegseis et al. 2006; Hawellek et al. 2013) that has been related to atrophy of the geniculocortical tracts (Shimony et al. 2006; Hawellek et al. 2013). The reduced alpha-band power, however, was not specific to working memory training in our study (Fig. S2 B).

### Working Memory Network Differences Prior to the Training

In our study, we used a working memory training paradigm to induce neuroplasticity and test differences in the formation of new networks in congenitally blind and sighted individuals. However, differences in working memory networks between the groups might have existed prior to the training due to long-term plasticity following early visual deprivation, and the training might have enhanced these differences. We tested this in an analysis where differences in theta-, beta- and gamma-band connectivity and power between blind and sighted participants in the pre-training 2-back session (contrasted with resting state data recorded at the beginning of the first MEG session) were compared. The analyses showed no significant effects, using the rigorous control for multiple comparisons. As these analyses were thought of as follow-up exploratory analyses, however, we performed another set of tests with a more liberal control for multiple-comparison of voxels at each ROI. The blind participants showed reduced theta-band connectivity compared to the sighted between the insula ROI and the IFG, the STG and pre- and post-central areas (Q = 0.2; FDR corrected α = .0298; p-values < .0298; Fig. S3 A). There were no differences at any other ROI (Q = 0.2; frontal: FDR corrected α = .0057; p-values > .0057; auditory: FDR corrected α = .0023; p-values > .0023; occipito-temporal and parietal: FDR corrected α = .0001; p-values > .0086) or frequency-band (beta-band: Q = 0.2; FDR corrected α=. 0001; frontal: p-values ≥ .0065; insula: p-values ≥ .0003; parietal: p-values ≥ .0004; temporal and occipito-temporal: p-values > .0001; gamma-band: Q = 0.2; FDR corrected α = .0001; frontal: p-values > .0001; insula: p-values ≥ .0026; parietal: p-values ≥ .0014; temporal: p-values ≥ .0037; occipito-temporal: p-values ≥ .0101) and no differences in power (beta- and gamma-band: Q = 0.2; FDR corrected α = .0001; beta-band: p-values ≥ .0007; gamma-band: p-values >.0001) (for between group differences in the resting state data: Fig. S3 B, C).

These findings suggest that differences in theta-band working memory networks between congenitally blind and sighted participants were partly present prior to the training, while differences in beta-band networks were largely established during the training.

## Discussion

The goal of this study was to extend previous reports on visual cortex activation during non-visual tasks in congenitally blind individuals by providing insights into the mechanisms underlying crossmodal neuroplasticity. We demonstrated a frequency-specific integration of visual cortex into task-specific networks in congenitally blind individuals during the performance of a working memory task with voices (Fig. 5). Working memory training with voices increased beta-band connectivity between visual cortex and brain areas associated with auditory working memory in congenitally blind adults. Visual cortex, furthermore, seems to be integrated in a functionally specific manner, particularly involving the right fusiform face area (Renier et al. 2014). Crucially, working memory training altered spectro-temporal characteristics of the networks differently in the blind (beta-band) and the sighted (theta-band), possibly reflecting a shift in the relative contribution of feedforward and feedback communication within the working memory network of blind individuals (Schmiedt et al. 2014; Bastos et al. 2015).

**Figure 5:**
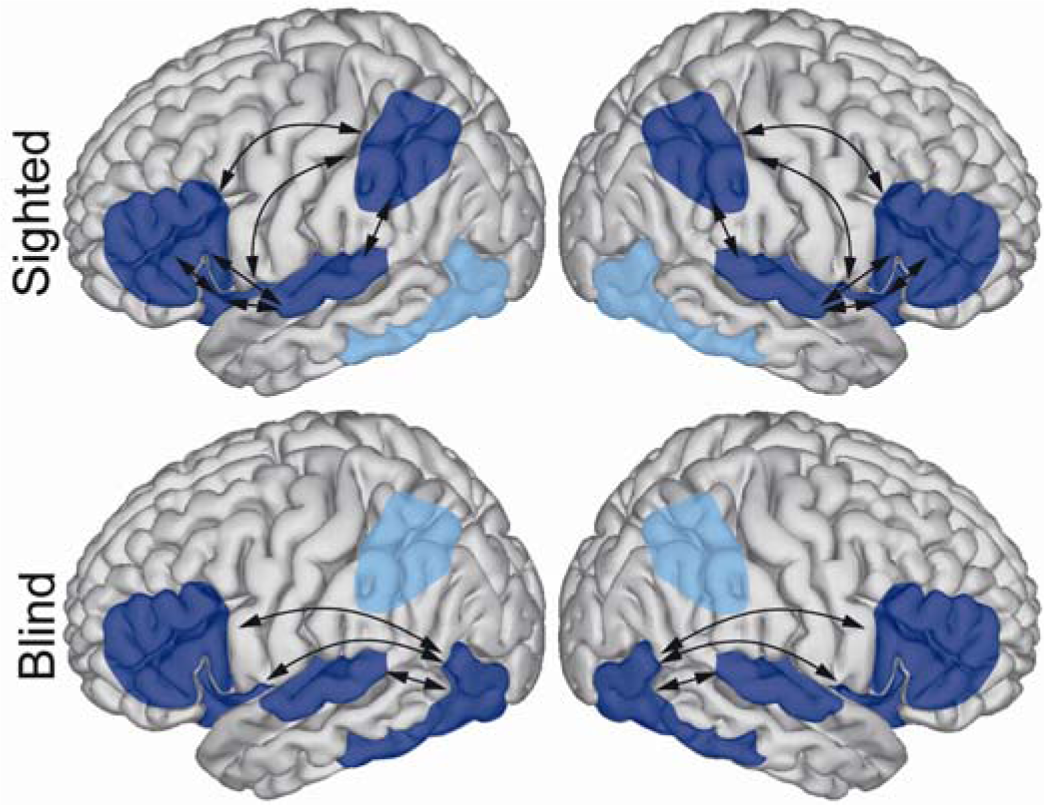
Schematic of the Networks Affected by Working Memory Training. *Connectivity increases due to working memory training were observed in the theta-band for sighted and in the beta-band for congenitally blind participants. All regions of interest (ROIs) that showed significant changes in connectivity to other brain areas (indicated by arrows) in the analysis of training-related differences between congenitally blind and sighted participants are displayed in dark blue. ROIs that did not show effects are shown in light blue.*

In both blind and sighted participants working memory training affected brain areas typically associated with auditory working memory processing, such as the ventrolateral prefrontal cortex (Plakke et al. 2015; Cohen et al. 2014; Plakke and Romanski 2014), the inferior parietal cortex (Jonides et al. 1998; Owen et al. 2005), the insula (Koelsch et al. 2009; Huang et al. 2013) and the auditory cortex (Bancroft et al. 2014). The inferior frontal gyrus, the superior temporal sulcus and the insular cortex have been previously shown to interact during voice identity processing (McGettigan et al. 2013), suggesting that in the present study blind and sighted participants recruited a working memory network specific to voice recognition. In the sighted participants working memory training with voices increased connectivity between these brain areas, possibly indicating a strengthening of or a higher processing efficiency within the task-relevant working memory network (Kelly and Garavan 2005).

Crucially, congenitally blind participants compared to the sighted showed a stronger training-related increase in connectivity between the visual cortex, including the right fusiform face area, and brain areas of the task-associated working memory network. The findings extend results from previous fMRI studies that described activation of fronto-parietal brain areas during working memory processing in congenitally blind and sighted individuals, while visual cortex was additionally activated in the blind (Amedi et al. 2003; Park et al. 2011; Deen et al. 2015). Note, that although the spatial resolution of source reconstruction in MEG is limited, our findings are in line with previous fMRI research. Previous fMRI studies found increased activation of the right fusiform face area (Hölig et al. 2014), an area particularly involved in the visual processing of faces (Kanwisher et al. 1997), as well as increased activation of the left STS and bilateral fusiform gyrus (Gougoux et al. 2009) in congenitally blind individuals during voice recognition. Both, voices and faces are highly familiar stimuli that convey human characteristics highly relevant for our everyday life. Crossmodal face-voice priming studies have shown that healthy sighted individuals are able to combine identity information from faces and voices (Ellis et al. 1997). Multisensory audiovisual integration areas, particularly involving the STS but additionally the posterior cingulate cortex, might provide *person identity nodes* (Campanella and Belin 2007). Furthermore, brain areas involved in multisensory processing could affect unisensory face and voice processing areas through top-down control, or direct coupling between unisensory areas (Campanella and Belin 2007). Evidence for a direct connection between unisensory areas comes from research that shows increased functional connectivity between face- (fusiform face area) and voice-selective areas (along anterior STS; Belin et al. 2000, 2004) for familiar compared to unfamiliar voices or when face-voice associations are learned (Kriegstein et al. 2005; Campanella and Belin 2007). Our finding of training-related increased connectivity between auditory and visual areas in the congenitally blind might indicate unmasking or strengthening of cross-modal connections that are only used under certain conditions in sighted humans, that is, when faces and voices are associated (Pascual-Leone et al. 2005). Actually, in the current study, in the sighted connectivity increased between small parts of the visual cortex and the frontal and insular cortex (Fig. 4A and B). A possibility is that the blindfolding of sighted participants in our study might have resulted in visual cortex recruitment (Schlaug et al. 2000; Merabet et al. 2008). Importantly, our findings show that working memory training resulted in a significantly higher increase of visual cortex connectivity in the congenitally blind compared to the sighted participants, which was specific to the working memory task with voices.

Furthermore, working memory training effects had different spectro-temporal profiles in congenitally blind (beta-band) and sighted (theta-band) participants, suggesting different dynamics of network communication. Both theta- and beta-band oscillations, supposedly, play a role in large-scale cortical connectivity (Varela et al. 2001). In the current study working memory training overall had more pronounced effects on large-scale connectivity compared to local activations reflected in oscillatory power. Overall, data from an exceptionally high number of congenitally blind individuals was included in this study (n = 27), however, note that although common for this type of investigation, a rather small number of participants was included in each condition (n = 9), possibly reducing the statistical power of our analyses, and also contributing to the lack of findings on local activations. Theta-band oscillations have been ascribed a role for the maintenance of multiple sequential items in working memory (Roux and Uhlhaas 2014). Theta-band connectivity recorded during resting state EEG has been shown to be altered by working memory training (Langer et al. 2013), while connectivity (degree centrality) decreased anteriorly (prefrontal, premotor, right enthorinal cortex) and increased posteriorly (parietal, right superior temporal gyrus, left inferior frontal gyrus, insular cortex). Similarly, beta-band oscillations have been suggested to have a particular role in working memory maintenance processes (Engel and Fries 2010; Kopell et al. 2011; Weiss and Mueller 2012), and fronto-parietal beta-band connectivity has been shown to be altered by working memory training (Astle et al. 2015). In the current study, we found no evidence that the differences in the spectro-temporal profile of the working memory networks in congenitally blind and sighted participants, relate to differences in working memory performance or learning behavior in the 2-back task. Another possibility is that congenitally blind individuals apply different working memory strategies, resulting in the different spectro-temporal profile of the networks. Crucially, in the current study, we found no differences in the reported strategies (Table S3; Supporting Information).

We propose that the different spectral profiles observed in sighted and blind individuals arose from changes in the functional architecture of working memory networks in the congenitally blind, due to the integration of visual cortex into existing networks. Whether, the changes in functional architecture are directly related to structural changes, that is, whether new connections were build or rather old connections were reconfigured to form new functional networks, requires further investigations. Importantly, our paradigm benefits from a within-subject normalization (analyses of changes across pre-post sessions), which reduces the impact of task unrelated anatomical differences between groups on our findings. One possibility is that the different spectro-temporal profile in the blind and the sighted reflect differences in the relative contribution of neural feedforward and feedback communication within the working memory networks. Feedback communication in the infragranular layers of visual cortex of macaque monkey has been shown to occur in the beta-band, while activity in the theta- and gamma-band was involved in feedforward communication in supragranular layers of visual cortex, as measured with LFPs (Schmiedt et al. 2014; Bastos et al. 2015) (for similar findings in human auditory cortex: Fontolan et al. 2014). Crucially, resecting a portion of V1 in macaque monkey resulted in a release of the beta-band suppression in V4, and thus, increased beta-band activity (LFP) during visual stimulation (Schmiedt et al. 2014). A possibility is that changes in the thalamo-cortical connections to V1 in congenitally blind (Ptito et al. 2008), results in increased beta-band activity, affecting the spectro-temporal profile of auditory working memory networks in the congenitally blind. Note that visual inspection suggests an antagonism between theta- and beta-band networks (Fig. 3), such that when working memory training resulted in an increased connectivity in one frequency band, it rather decreased in the control group while connectivity in the other frequency band showed the opposite pattern across training groups. This pattern of increased and decreased connectivity differed between the congenitally blind and sighted group. This dissociation suggests that working memory training resulted at the same time in enhancement and suppression of distinct (theta- and beta-band) networks in the congenitally blind and sighted individuals. Further research, however, is required to better understand this effect.

In conclusion, our study provides novel evidence that congenital visual deprivation alters large-scale interactions of working memory related neural networks. In congenitally blind participants, parts of visual cortex were integrated into an auditory working memory network following a working memory training with auditory stimuli. Different spectral characteristics of networks in congenitally blind compared to sighted adults might indicate different neural coupling mechanisms, reflecting a change in the distribution of feedforward and feedback communication due to the integration of visual cortex into functional networks in the blind.

## Abbreviations

CRR: correct rejection rate
Cs: cross-spectra
FDR: False discovery rate
FFA: fusiform face area
fMRI: functional magnetic resonance imaging
hMT+/V5: occipital-temporal-parietal junction (human correlate of macaque middle temporal visual area)
HR: hitrate
IFG: inferior frontal gyrus
HSWBS: Habituelle Subjektive Wohlbefindens Skala
IPC: inferior parietal cortex
IPG: inferior parietal gyrus
IPL: inferior parietal lobe
ITI: inter-trial interval
ITL: inferior temporal lobe
LFP: local field potential
MEG: magnetoencephalography
MFG: middle frontal gyrus
MNI: Montreal Neurological Institute
MTG: middle temporal gyrus
Pow: power
PSQ-20: Perceived Stress Questionnaire
ROI: regions of interest
SD: standard deviation
STG: right superior temporal gyrus
STS: anterior superior temporal sulcus
VLMT: Verbaler Lern-und Merkfähigkeitstest.

## Acknowledgements

We want to thank Malte Sengelmann for technical support, Till R. Schneider and Georgios Michalareas for methodological advice and Mirja Biewener, and Constanze Mahnert for assistance with the experiment conduction.

